# Alzheimer’s disease BIN1 coding variants increase intracellular Aβ by interfering with BACE1 recycling

**DOI:** 10.1101/2021.06.09.447716

**Authors:** Catarina Perdigão, Mariana Barata, Tatiana Burrinha, Cláudia Guimas Almeida

## Abstract

Genetics identified *BIN1* as the second most important risk locus associated with late-onset Alzheimer’s disease, after APOE4. Here we show the consequences of two coding variants in BIN1 (rs754834233 and rs138047593), both in terms of intracellular beta-amyloid accumulation (iAbeta) and early endosome enlargement, two interrelated early cytopathological Alzheimer’s disease phenotypes, supporting their association with LOAD risk. We previously found that Bin1 deficiency potentiates beta-amyloid production by decreasing BACE1 recycling and enlarging early endosomes. Here, we demonstrate that the expression of the two LOAD mutant forms of Bin1 did not rescue the iAbeta accumulation and early endosome enlargement induced by Bin1 knockdown and recovered by wild-type Bin1. The LOAD coding variants reduced Bin1 interaction with BACE1 likely causing a dominant-negative effect since Bin1 mutants, but not wild-type Bin1, overexpression increased iAbeta42 due to defective BACE1 recycling and accumulation in early endosomes. Endocytic recycling of transferrin was similarly affected by Bin1 wild-type and mutants, indicating that Bin1 is a general regulator of endocytic recycling. These data show that the LOAD mutations in Bin1 lead to a loss of function, suggesting that endocytic recycling defects are an early causal mechanism of Alzheimer’s disease.

## INTRODUCTION

Alzheimer’s disease (AD) is the most common neurodegenerative disease worldwide. The earliest known mechanisms driving AD predicted to begin decades before diagnoses are beta-amyloid (Aβ) intracellular accumulation and endosome dysfunction (1, 2). Aβ is generated intracellularly through the sequential processing of the transmembrane amyloid precursor protein (APP) by β-secretase (BACE1) and γ-secretase (3–6). APP cleavage by BACE1 is the rate-limiting step to generate Aβ (4). Interestingly, APP and BACE1 segregate in the plasma membrane (7, 8). Endocytosis potentiates the encounter of APP and BACE1, and processing, by delivery to a common early endosome (7, 9–12). Aβ production is counteracted by APP sorting for degradation (13–15) and BACE1 recycling to the plasma membrane (8, 15). Endosomal dysfunction causes are unclear. In familial AD (FAD), autosomal mutations cause increased Aβ42 production (16–19), which we and others implicated in endosomal abnormalities (20–23). In LOAD, the causal mechanisms of Aβ intracellular accumulation and endosomal enlargement are likely different. LOAD is expected multifactorial, caused by a combination of aging, lifestyle, and genetic risk factors. Given the prediction for a strong genetic predisposition, 58 to 79% (24), geneticists have been looking for genetic risk factors in LOAD patients. Among the genetic risk factors identified by several genome-wide association studies (GWAS), *BIN1*, bridging integrator 1, was the second most associated with increased AD risk (25– 31).

*BIN1* encodes several isoforms and, in the brain, are mainly expressed the neuronal and ubiquitous isoforms (32). Bin1 belongs to the BAR (Bin1/amphiphysin/RVS167) superfamily. Bin1 isoforms share the N-BAR domain, responsible for sensing and inducing curvature of membranes (33, 34), and the SH3 domain, responsible for interacting with several endocytic players, such as dynamin (35–38), involved in the scission of budding vesicles. The neuronal-specific isoform also encodes the CLAP (clathrin and AP2 binding) domain, responsible for interacting with clathrin and AP2 (39), both required for clathrin-mediated endocytosis. In non-neuronal cells, Bin1 overexpression inhibits transferrin endocytosis, known to be mediated by clathrin (38). Furthermore, Bin1 knockdown reduces transferrin receptor recycling but not its endocytosis (40, 41). In neurons, we previously showed that Bin1 polarizes to axons, associated with early endosomes (15).

In AD, how Bin1 levels change is still controversial. *BIN1* transcripts increase in AD human brains (42). However, lower *BIN1* transcripts correlate with earlier disease onset (43). In FAD models, Bin1 protein accumulates adjacent to amyloid plaques (44). In contrast, in LOAD human brain homogenates, Bin1 protein levels decrease (45, 46) or are unchanged (47). An analysis of Bin1 isoforms separately revealed that neuronal Bin1 decreases while ubiquitous Bin1 increases in AD human brains (48, 49).

To study the impact of Bin1 depletion, researchers have taken a knockdown approach *in vitro* because the Bin1 mouse knockout is perinatal lethal (40). Bin1 knockdown in cortical neurons increases Aβ42 intracellular production (15, 50). In addition, Bin1 knockdown reduces endocytic BACE1 recycling (15), probably enlarging early endosomes (15, 51). Mechanistically, Bin1 contributes to the scission of recycling carriers containing BACE1 from early endosomes (15). *In vivo* Aβ accumulation was undetectable in mice conditionally knocked-out for Bin1 in excitatory neurons (52), indicating that Bin1 does not control the Aβ production in excitatory neurons or the intracellular Aβ accumulation is difficult to detect *in vivo* (53). These findings support Bin1 loss of function in AD, implicated in AD earliest mechanisms in neurons: Aβ intracellular accumulation and endosomal abnormalities.

The impact of Bin1 accumulation in AD is less studied. Increased Bin1 expression decreases early endosomes size (51), opposite to AD early endosome enlargement but possibly linked to tau spreading, a mechanism related to AD progression. However, whether Bin1 increased levels impact Aβ42 intracellular accumulation is still not known.

GWAS and subsequent targeted sequencing associated *BIN1* variants, in regulatory and coding regions, with LOAD and poorer memory performance (25–30, 32, 54, 55). While the regulatory variants may be more frequent and likely associated with alterations in Bin1 transcription, the impact of the coding variants in Bin1 is unknown. Two coding variants leading to mutations in Bin1 were associated with LOAD (54, 56). The first identified was rs754834233 (P318L (PL)), a proline for a leucine mutation localized to the proline-serine-rich domain proximal to the CLAP domain (56). The second mutation identified was rs138047593 (K358R (KR)), an arginine for a lysine mutation within the Bin1 SH3 domain (54). Both mutations locate in or near domains necessary for Bin1 proper function at endocytosis and recycling.

We set out to investigate if two LOAD Bin1 mutations interfere with Bin1 function and lead to LOAD earliest mechanisms, Aβ intracellular accumulation, and endosomal abnormalities. We mutagenized wild-type Bin1 with LOAD Bin1 PL and KR mutations. We used an overexpression and rescue approach in the neuronal N2a cell line. By analyzing endogenous intracellular Aβ42 accumulation, BACE1 endocytic trafficking, and early endosome size, we found that PL and KR replicate the impact of Bin1 loss of function. Thus, these mutations may contribute to the development of early mechanisms of LOAD.

## RESULTS

### Bin1 mutants increase intracellular Aβ42 accumulation

Previously, we found that Bin1 loss of function results in Aβ42 intracellular accumulation in a neuroblastoma cell line (N2a) and murine primary neurons (15). Significantly, this defect was only rescued by Bin1 neuronal isoform (15), revealing a specific function of neuronal Bin1 in Aβ42 production. We now want to understand if rare coding variants that lead to mutations in Bin1 found associated with LOAD are sufficient to increase Aβ42 intracellular accumulation. Besides, we investigate if neuronal Bin1 increased levels also interfere with Aβ42 accumulation. To do so, we used a semi-quantitative assay for intracellular endogenous Aβ based on Aβ42 immunofluorescence that we performed previously (15, 20, 53). We have controlled this assay extensively, including with APP knockout cells (57). Here, we show that Aβ42 immunofluorescence was significantly reduced (50%) upon inhibition of Aβ production (Fig. S1 A and B). We introduced PL and KR mutations, in the corresponding nucleotides, in mouse neuronal MYC-tagged BIN1 cDNA (Bin1-PL and Bin1-KR, respectively). We overexpressed (OE) either neuronal Bin1 wild-type (Bin1-WT), Bin1-PL, Bin1-KR, or MYC-empty vector (MYC) in N2a cells. We observed that overexpressed Bin1-PL and KR have a broad cellular distribution similar to overexpressed Bin1-WT (Fig. 1A) but different from endogenous Bin1, which localizes to early endosomes(15). We found that Bin1-WT overexpression did not change intracellular endogenous Aβ42 levels (Fig. 1A and B). In contrast, Bin1-PL and Bin1-KR overexpression increased intracellular Aβ42 significantly, in 12% and 30%, respectively. Since the fluorescence of Aβ42 increased in areas of high anti-MYC fluorescence, we analyzed at higher resolution the fluorescence profiles of Aβ42, Bin1-WT, Bin1 mutants, or MYC vector and found that the fluorescence peaks do not entirely overlap (Fig. S1C and D). Moreover, we verified that the anti-MYC immunofluorescence mean intensity was unchanged when Aβ42 mean intensity was reduced (Fig. S1A and B).

**Figure 1.**
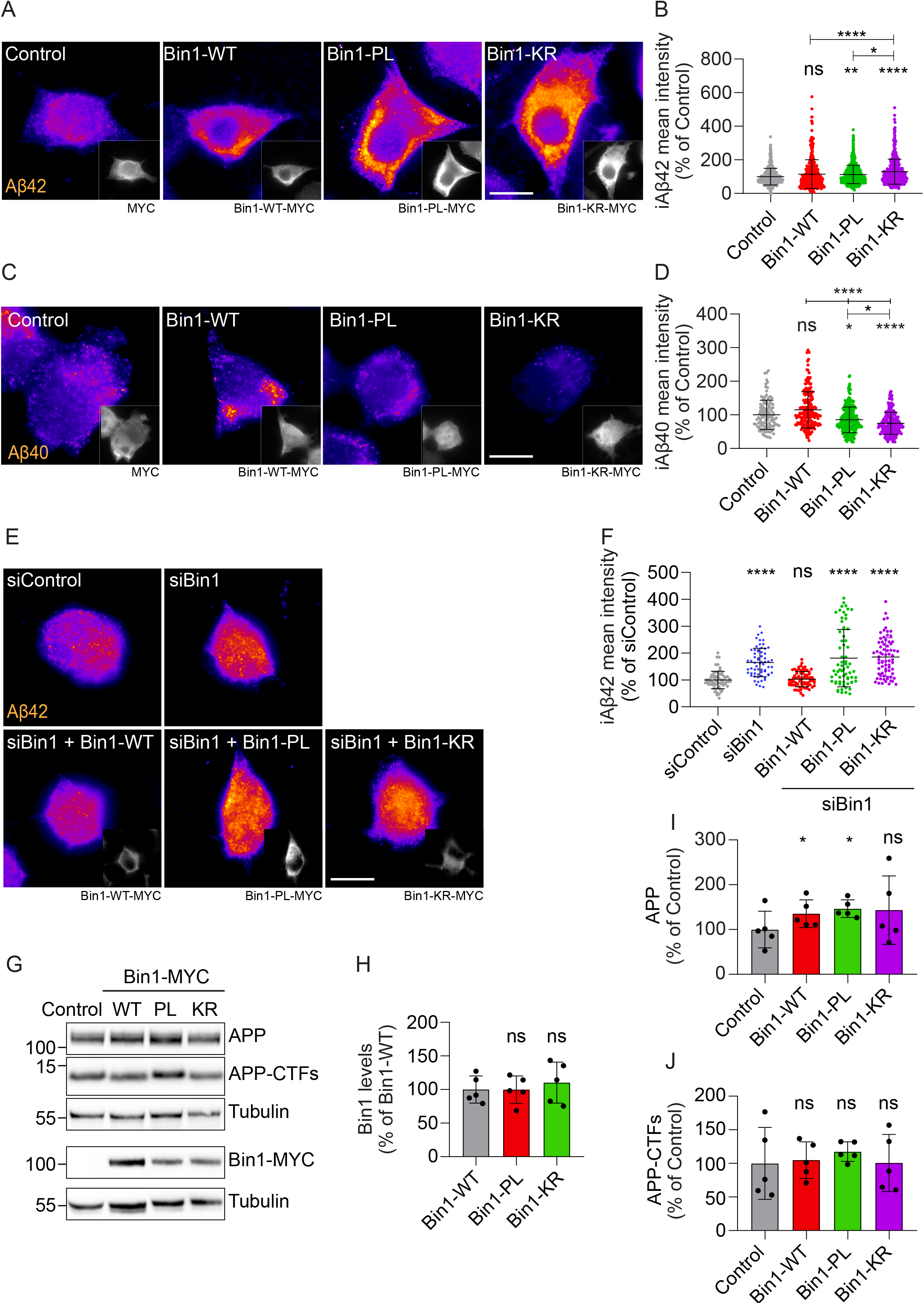
BIN1 mutants increase intracellular Aβ accumulation. **(A)** Intracellular endogenous Aβ42 (fire LUT) in N2a cells transiently expressing Bin1 wild-type (WT), Bin1 mutants PL and KR tagged with MYC, or MYC (control), immunolabelled with anti-Aβ42 and anti-MYC (insets), analyzed by epifluorescence microscopy. Scale bar, 10 µm. **(B)** Quantification of mean Aβ42 fluorescence intensity relative to control cells (n=4, N_control_=492 cells, N_Bin1-WT_=410 cells, N_Bin1-PL_=564 cells, N_Bin1-KR_=530 cells; \**P*=0.0284 Bin1-PL vs. Bin1-KR, \*\**P*=0.0018 Bin1-PL vs. control, \*\*\*\**P*<0.0001 Bin1-WT vs. Bin1-KR, Bin1-KR vs. control, Kruskal-Wallis test, mean ± SD). **(C)** Intracellular endogenous Aβ40 (fire LUT) in N2a cells transiently expressing Bin1 wild-type (WT), Bin1 mutants PL and KR tagged with MYC, or MYC (control), immunolabelled with anti-Aβ40 and anti-MYC (insets), analyzed by epifluorescence microscopy. Scale bar, 10 µm. **(D)** Quantification of mean Aβ40 fluorescence intensity relative to control cells (n=3, N_control_=114 cells, N_Bin1-WT_=187 cells, N_Bin1-PL_=251 cells, N_Bin1-KR_=224 cells; ^ns^*P*=0.2287 Bin1-WT vs. control, \**P*=0.0333 Bin1-PL vs. control, ****P<0.0001 Bin1-WT vs. control, ****P<0.0001 Bin1-PL vs Bin1 - WT, ****P<0.0001 Bin1-KR vs. Bin1-WT, \**P*=0.0333 Bin1-KR vs. Bin1-PL, Kruskal-Wallis test, mean ± SD). **(E)** Intracellular endogenous Aβ42 (fire) in siControl- and siBin1-treated N2a cells followed by transient transfection of BIN1-WT or BIN1 mutants PL and KR tagged with MYC. N2a cells immunolabelled with anti-Aβ42 (upper panel) and anti-MYC (lower panel), analyzed by epifluorescence microscopy. Scale bar, 10 μm. **(F)** Quantification of intracellular Aβ42 (iAβ42) mean fluorescence in percentage of siControl (n=3, N_siControl_=64 cells, N_siBin1_=63 cells, N_siBin1+Bin1-WT_=65 cells, N_siBin1+Bin1-PL_=67 cells, N_siBin1+Bin1-KR_=72 cells; \*\*\*\**P*<0.0001 siBin1 vs. siControl, siBin1+Bin1-PL vs. siControl, siBin1+Bin1-KR vs. siControl, Kruskal-Wallis test, mean ± SD). **(G)** Endogenous APP and APP-CTFs levels by western blot with anti-APP antibody (Y188) of N2a cells transiently expressing Bin1-WT, or BIN1-PL and -KR tagged with MYC, or MYC (control). Bin1 expression was analyzed by western blot with anti-MYC antibody. MYC empty vector was not detectable due to its small size. Tubulin was immunoblotted as the loading control. **(H)** Quantification of Bin1-WT, Bin1-PL, and Bin1-KR levels normalized to tubulin in the percentage of Bin1-WT (n=5, ^ns^*P*=0.8649 Bin1-KR vs. Bin1-WT, ^ns^P=0.9829 Bin1-PL vs.Bin1-WT, RM one way-ANOVA, mean ± SD). **(I)** Quantification of APP levels normalized to tubulin in percentage of control (n=5, ^ns^*P*= 0.2236 Bin1-KR vs. control, \**P*=0.0227 Bin1-PL vs. control, \**P*=0.0171 Bin1-WT vs. control, paired t-test, mean ± SD). **(J)** Quantification of APP-CTFs levels normalized to tubulin in the percentage of control (n=5, ^ns^*P*=0.9831 Bin1-PL vs. control, ^ns^*P*=0.9965 Bin1-WT vs. control, Bin1-KR vs. control, RM one way-ANOVA, mean ± SD).

To understand whether Bin1 mutants induced Aβ42 accumulation in early or late endosomes/lysosomes, we assessed Aβ42 colocalization with EEA1- or LAMP1-positive endosomes, the Bin1-PL only increased Aβ42 in LAMP1-positive endosomes (Fig. S2A and B). Overall, the data respectively. We found that Bin1-KR increased Aβ42 in EEA1- and LAMP1-positive endosomes, while indicate that Bin1 mutants increase Aβ42 more in late-endosomes/lysosomes (40%) than in early endosomes (30%). This trend is in agreement with our previous observations in primary neurons modeling familial AD (21).

To confirm if the rise in Aβ accumulation was specific for Aβ42, the most toxic Aβ, we evaluated the effect of Bin1-WT and mutants’ overexpression in intracellular Aβ40 (Fig. 1C and D). Indeed, we more Aβ40 (25%), in contrast to the observed increase in Aβ42. Of note, Bin1-WT overexpression found that Bin1-PL overexpression decreased Aβ40 (15%), and the Bin1-KR overexpression reduced showed no significant impact on Aβ40 levels (25%). Together these results suggest that Bin1 mutants may contribute to an increased ratio of Aβ42 over 40, potentially linked to higher Aβ toxicity (58).

The increase in Aβ42 accumulation upon Bin1 mutants’ overexpression recapitulates the impact of Bin1 knockdown (KD) (15, 50), suggesting that the PL and KR mutations lead to a Bin1 loss of function with a dominant-negative effect over the endogenous Bin1.

To verify if these mutations lead to a loss of function, we eliminated the confounding role of endogenous Bin1 by expressing siRNA-resistant Bin1-WT or Bin1 mutants in cells treated with Bin1 siRNA (knockdown). The localization of Bin1 mutants was similar to Bin1-WT when expressed in Bin1 knockdown (KD) cells (Fig. 1E). Importantly, we observed that the mutants did not rescue the rise in Aβ42 levels induced Bin1 KD, as the neuronal Bin1-WT (15) (Fig. 1E and F). This result supports that the LOAD mutations cause a loss of function of Bin1 in the control of Aβ production.

To understand if increased APP processing underscored the rise in Aβ42, we investigated APP processing into its C-terminal fragments (APP-CTFs). APP processing was analyzed by western blot using the antibody Y188 against the C-terminal domain of APP, detecting APP full-length and the APP-CTFs. We did not see changes in the level of endogenous APP-CTFs when expressing Bin1-WT or its mutants (Fig. 1G and J). Instead, we found APP full-length increased upon Bin1-WT (35%) and Bin1-PL (46%) but not Bin-KR overexpression (Fig. 1I). This increase in APP full-length does not correlate with the observed rise in Aβ levels. While Aβ is higher with the KR mutant, APP full-length is not. We also observed that Bin1 mutants did not change Bin1 expression levels (Fig. 1G and H). Since Bin1-WT OE decreases early endosomes (51), it could decrease sorting at early endosomes for lysosomal degradation, explaining the increase in APP levels.

### Bin1 mutants lose control of early endosomes size

Next, we investigated if the LOAD mutations in Bin1 lead to endosomal abnormalities, namely endosomal enlargement, another early LOAD mechanism. Previous work demonstrated that Bin1 controls early endosome size since Bin1 KD increases it and neuronal Bin1 OE reduces it (15, 51).

To investigate how Bin1 mutants affect early endosome size, we transiently expressed Rab5-GFP, a RAB GTPase enriched at early endosomes, overexpressed Bin1-WT, Bin1-PL, Bin1-KR, and MYC as control, and measured Rab5-positive endosome size in N2a cells. As previously reported, Bin1-WT OE decreased Rab5-positive endosome size by 20% (Fig. 2A and B). Differently, the Bin1-PL OE reduced Rab5-positive endosome size only by 10%, and the Bin1-KR OE did not alter endosome size (Fig. 2A and B). Additionally, we analyzed endogenous EEA1, another marker of early endosomes (Fig.S1). Similarly to Rab5, we found a shrinkage of EEA1-positive endosomes upon Bin1-WT OE (18%). The PL and KR mutants induced smaller reductions in EEA1-positive endosome size, by 13% and 4%, respectively, although only the KR mutant was significantly different from Bin1-WT (Fig.S1 A and B). Of note, Bin1-WT OE led to an increase in the number of Rab5- and EEA1-positive endosomes by 22% and 24%, respectively (Fig.S1 C and D). The Bin1 mutants failed to increase the number of EEA1-positive endosomes but not Rab5-positive endosomes (Fig.S1 C and D), suggesting that the Rab5 OE compensates for the Bin1 mutants’ effect on endosomes.

**Figure 2.**
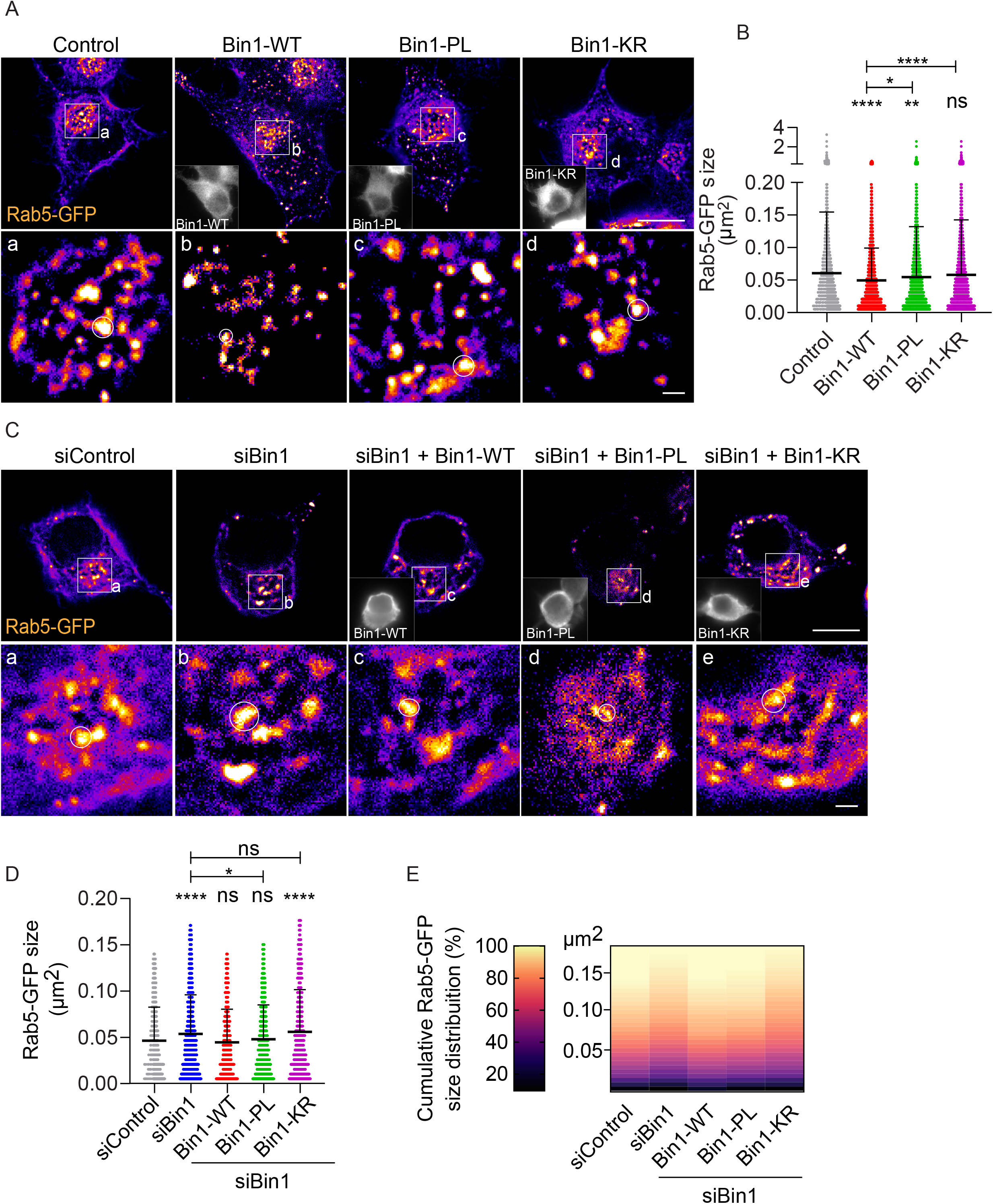
Bin1 mutants lose control of early endosomes size. **(A)** Bin1 mutants’ overexpression impact early endosomes. Rab5-GFP positive early endosomes (fire LUT) detected in N2a cells transiently expressing Rab5-GFP (control) and Bin1-WT or Bin1-PL and - KR tagged with MYC (insets), analyzed by epifluorescence microscopy. Images are displayed after background subtraction with Fiji. Scale bar, 10 μm. The white squares indicate the perinuclear region magnified in (a-d), where the white circles highlight Rab5-positive early endosomes mean size. Scale bar, 1 μm. **(B)** Quantification of Rab5-positive early endosomes size (µm^2^). (n=3, N_control_=96 cells, N_Bin1-WT_=45 cells, N_Bin1-PL_=51 cells, N_Bin1-KR_=54 cells; ^ns^*P*=0.1409 Bin1-KR vs. control, \**P*=0.0300 Bin1-WT vs. Bin1-PL, \*\**P*=0.0027 Bin1-PL vs. Control, \*\*\*\**P*<0.0001 Bin1-WT vs. control, Bin1 WT vs. Bin1 KR, one way-ANOVA with Tukey’s multiple comparisons test, mean ± SD). **(C)** Bin1 mutants rescue of early endosomes enlargement induced by Bin1 KD. Rab5-positive early endosomes (fire LUT) were detected in siControl- and siBin1-treated N2a cells alone or upon transient transfection with Bin1-WT Bin1-PL and -KR tagged with MYC (insets), analyzed by epifluorescence microscopy. Images are displayed after background subtraction with Fiji. Scale bar, 10 μm. The white squares indicate the perinuclear region magnified in (a-d), where the white circles highlight Rab5-positive early endosomes’ mean size. Scale bar, 1 μm. **(D)** Quantification of Rab5-positive early endosomes size (µm^2^) (n=3, N_siControl_=48 cells, N_siBin1_=51 cells, N_siBin1+Bin1-WT_=31 cells, N_siBin+Bin1-PL_=39 cells, N_siBin1+Bin1-KR_=36 cells; ^ns^*P*>0.9999 siBin1+Bin1-WT vs. siControl, siBin1+Bin1-PL vs. siControl, siBin1 vs siBin1+Bin1 KR, \**P*=0.0243 siBin1 vs. siBin1+Bin1-PL, \*\*\*\**P*<0.0001 siBin1 vs. siControl, siBin1+Bin1-KR vs. siControl, Kruskal-Wallis test, mean ± SD). **(E)** Cumulative Rab5-positive endosomes size (µm^2^) frequency distribution. Colormap magma: 0% (black) to 100% (yellow) (n=3; N_siControl_=48 cells, N_siBin1_=51 cells, N_siBin1+Bin1-WT_=31 cells, N_siBin+Bin1-PL_=39 cells, N_siBin1+Bin1-KR_=36 cells).

To remove the confounding role of endogenous Bin1, we performed a rescue experiment in which we expressed Bin1-WT and mutants upon Bin1 KD with siRNA. As reported, we found Rab5-positive endosomes 16% larger upon Bin1 KD (15, 51). Furthermore, Bin1-WT re-expression rescued Rab5-positive endosomes size, whereas Bin1-PL rescued partially, and Bin1-KR did not rescue (Fig. 2C and D). Additionally, a heatmap of cumulative distribution (%) of endosome size is shown (Fig. 2F). The deepening of greater size blocks for siBin1 indicates the higher percentage of larger endosomes, similar after Bin1-KR re-expression but different from Bin1-WT and Bin1-PL re-expression.

These results suggest that the KR mutation is more pathogenic than the PL mutation, inducing a complete Bin1 loss of function in the control of early endosomes size.

### Bin1 mutations reduce interaction with BACE1 and its endocytic recycling

Previously, we linked the increase in endosome size to the decreased recycling of BACE1 when Bin1 was KD (1). In addition, Miyagawa et al. showed that Bin1 interacts with BACE1 in HeLa cells (50). Here, we demonstrate that Bin1 co-immunoprecipitated BACE1 from wild-type mouse brain lysates (Fig. 3A), supporting that Bin1 interacts with BACE1 in the brain. Moreover, we show the reverse that BACE1-GFP co-immunoprecipitated Bin1-WT from N2a cells co-expressing Bace1-GFP and Bin1-WT (Fig. 3B). We assessed if the LOAD mutations interfere with the Bin1 interaction with BACE1. Importantly, BACE1-GFP co-immunoprecipitated less Bin1-PL and Bin1-KR (Fig. 3B). Quantification showed that the BACE1-GFP tended to interact less with Bin1-PL while the interaction with Bin1-KR was significantly reduced (65%) (Fig. 3C), which indicates that although both mutations may interfere with Bin1 interaction with BACE1, the KR mutation has a more disruptive effect.

**Figure 3.**
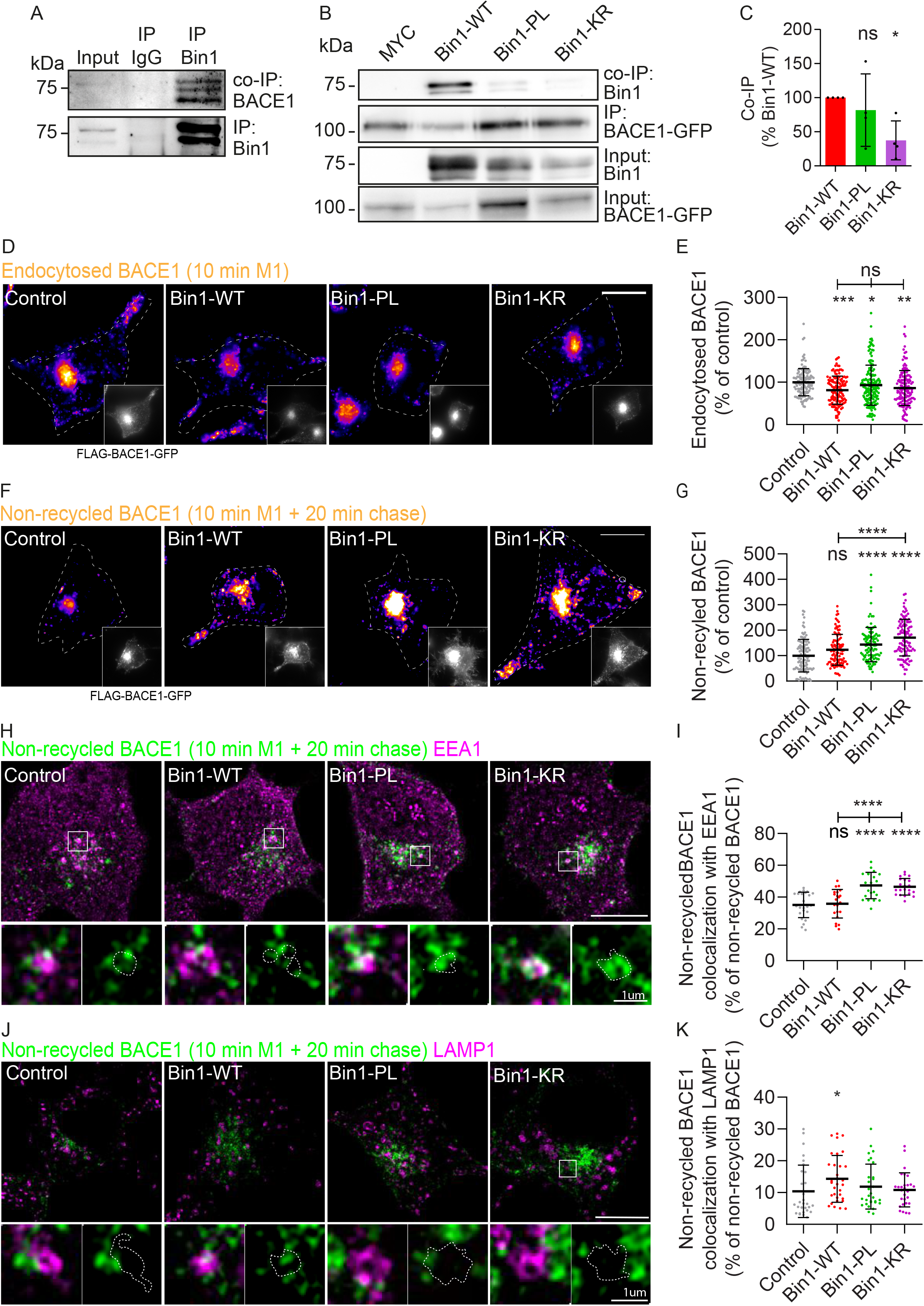
Bin1mutants’ impact BACE1 endocytic trafficking. **(A)** BACE1 co-immunoprecipitation with Bin1 from wild-type mouse brain homogenates (input) using anti-Bin1 (IP Bin1), or normal IgG (IP IgG) detected with anti-BACE1 or anti-Bin1 antibodies (n=3). **(B)** Bin1-WT or mutants co-immunoprecipitation with BACE1-GFP from cells co-expressing BACE1-GFP and Bin1-WT, Bin1-PL or Bin1-KR (input), using GFP-traps, detected with anti-BACE1 (IP) or anti-Bin1 (Co-IP) antibodies. **(C)** Quantification of co-immunoprecipitated Bin1-WT or mutants normalized by immunoprecipitated BACE1 and respective inputs (n=4, ^ns^*P*=0.3143 Bin1-PL vs. Bin1-WT, ^ns^*P*=0.3429 Bin1-PL vs. Bin1-KR, \**P*=0.0286 Bin1-WT vs. Bin1-KR. (**D-K**) BACE1 endocytic trafficking followed in N2a cells transiently expressing BACE-GFP N-terminally tagged with FLAG (control) and BIN1-WT or BIN1-PL and -KR, using a pulse-chase assay with M1, an anti-FLAG antibody, analyzed by epifluorescence microscopy. **(D)** Endocytosed BACE1 (10 min M1, fire LUT) assessed by immunofluorescence in N2a cells with a secondary antibody against the endocytosed anti-FLAG (M1). Insets show GFP signal corresponding to FLAG-BACE1-GFP. Scale bar, 10 μm. **(E)** Quantification of endocytosed BACE1 (10 min M1) fluorescence normalized to FLAG-BACE1-GFP fluorescence and in the percentage of the control (n=4, N_control_=134 cells, N_Bin1-WT_=137 cells, N_Bin1-PL_=171 cells, N_Bin1-KR_=142 cells, ^ns^*P*=0.5282 Bin1-WT vs. Bin1-PL, ^ns^*P*>0.9999 Bin1-WT vs. Bin1-KR, Bin1-PL vs. Bin1-KR, \**P*=0.0404 Bin1-PL vs. control, \*\**P*=0.0017 Bin1-KR vs. control, \*\*\**P*=0.0002 Bin1-WT vs. control, Kruskal-Wallis test, mean ± SD). **(F)** Non-recycled BACE1 (fire LUT) detected upon 10 min pulse with M1 and 20 min chase, assessed by immunofluorescence in N2a cells with a secondary antibody against the endocytosed anti-FLAG (M1). Insets show GFP signal corresponding to FLAG-BACE1-GFP. Scale bar, 10 μm. **(G)** Quantification of non-recycled BACE1 fluorescence normalized to FLAG-BACE1-GFP fluorescence and in the percentage of the control (n=3, N_control_=105 cells, N_Bin1-WT_=101, N_Bin1-PL_=104 cells, N_Bin1-KR_=104 cells; ^ns^*P*=0.0934 Bin1-WT vs. Control, \*\*\*\**P*<0.0001 Bin1-PL vs. control, Bin1-KR vs. control, Bin1-WT vs. Bin1-KR, Kruskal-Wallis test, mean ± SD). **(H)** Non-recycled BACE1 (green, detected upon 10 min pulse with M1 and 20 min chase) localization in EEA1-positive endosomes (magenta), assessed by immunofluorescence in N2a cells with an antibody against endogenous EEA1. Images are displayed merged after background subtraction with Fiji. Scale bar, 10 μm. The white squares indicate magnified endosomes. Non-recycled BACE1 is shown individually and merged with EEA1. Scale bar, 1 μm. **(I)** Quantification of non-recycled BACE1 colocalization with EEA1 (n=3, N_control_=24 cells, N_Bin1-WT_=22 cells, N_Bin1-PL_=21 cells, N_Bin1-KR_=21 cells; ^ns^*P=*0.9305 Bin1-WT vs. control, \*\*\*\**P*<0.0001 Bin1-PL vs. control, Bin1-KR vs. control, Bin1-WT vs. Bin1-PL, Bin1-WT vs. Bin1-KR, one-way ANOVA, mean ± SD). **(J)** Non-recycled BACE1 (green, detected upon 10 min pulse with M1 and 20 min chase) localization in LAMP1-positive endosomes (magenta), assessed by immunofluorescence in N2a cells with an antibody against endogenous LAMP1. Images are displayed merged after background subtraction with Fiji. Scale bar, 10 μm. The white squares indicate magnified endosomes. Non-recycled BACE1 is shown individually and merged with LAMP1. Scale bar, 1 μm. **(K)** Quantification of non-recycled BACE1 colocalization with LAMP1 (n=3, N_control_=29 cells, N_Bin1-WT_=32 cells, N_Bin1-PL_=30 cells, N_Bin1-KR_=28 cells; ^ns^*P>*0.9999 Bin1-KR vs. control, Bin1-WT vs. Bin1-PL, Bin1-PL vs. Bin1-KR, ^ns^*P=*0.8539 Bin1-PL vs. control, ^ns^*P=*0.6546 Bin1-WT vs. Bin1-KR, \**P=*0.0299 Bin1-WT vs. control, Kruskal-Wallis test, mean ± SD).

BACE1 trafficking could be affected by the loss of its interaction with Bin1 mutants. To confirm this hypothesis, we investigated if the LOAD mutations alter Bin1 control of BACE1 endocytic recycling. To analyze BACE1 trafficking, we performed pulse/chase assays using an antibody against FLAG (M1) in N2a cells transiently expressing BACE1 cDNA with an N-terminal FLAG-tag and a C-terminal GFP (FLAG-BACE1-GFP), as previously (15).

To measure BACE1 recycling to the plasma membrane, we pulsed N2a cells with M1 for 10 min, then we acid-stripped non-endocytosed BACE1-bound M1 and further chased endocytosed BACE1-bound M1 for 20 min (15, 59).

Firstly, we measured BACE1 endocytosis (10 min pulse) since Bin1-WT OE decreases transferrin endocytosis (60), which we confirmed (Fig. 4). Endocytosed BACE1 was delivered to EEA1-positive early endosomes (Fig. S4A). We found that Bin1-WT OE reduced BACE1 endocytosis by 15% (Fig. 3D and E). The mutations in Bin1 did not alter the BACE1 endocytosis decrease induced by Bin1-WT OE (10% Bin1-PL and 15% Bin1-KR; Fig. 3D and E). Bin1-WT likely alters endocytosis when overexpressed by sequestering necessary endocytic components since Bin1 is not required for BACE1 endocytosis (15). The LOAD mutations do not change this overexpression phenotype.

**Figure 4.**
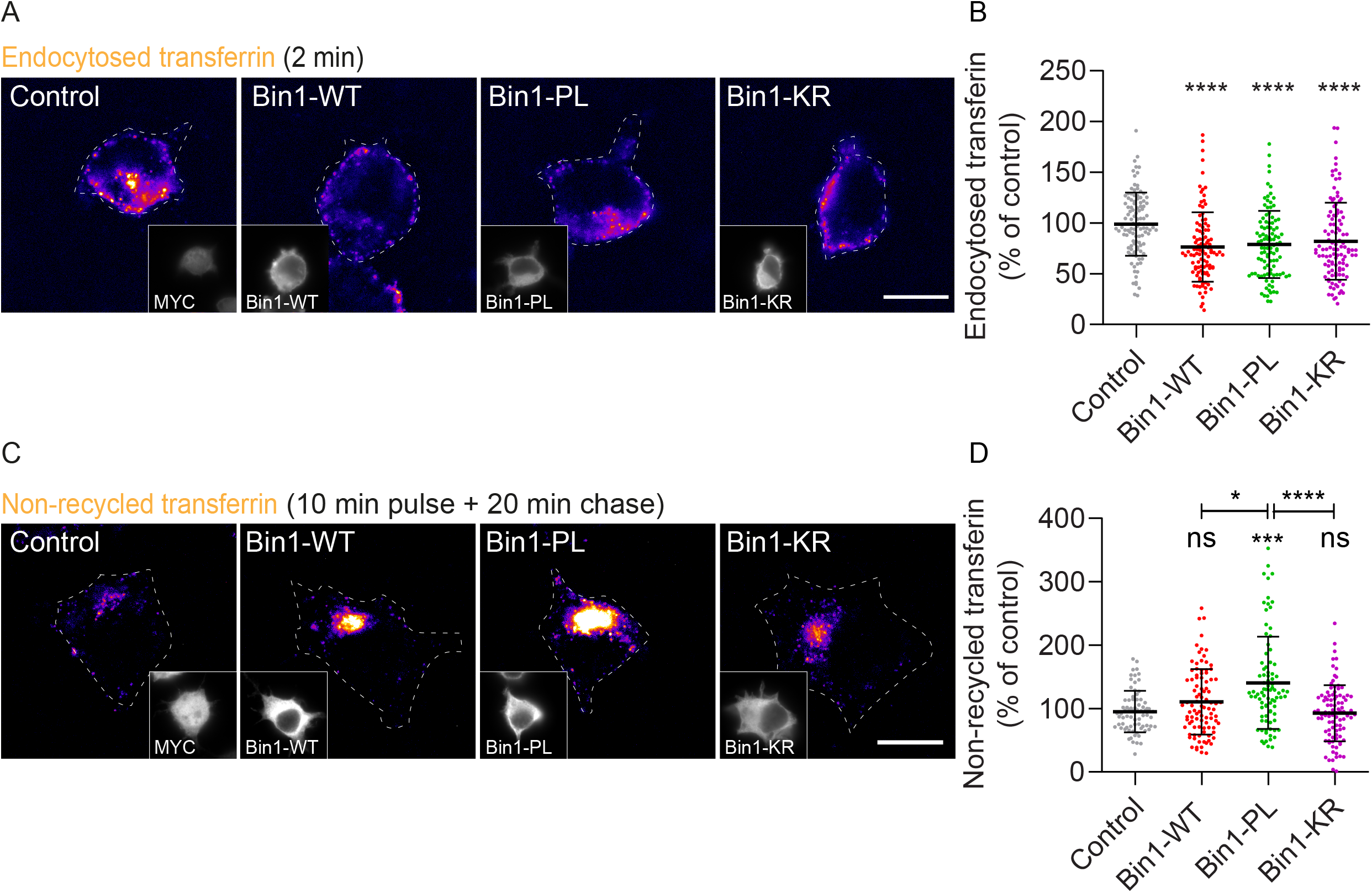
Bin1 mutants impair the canonical transferrin endocytic recycling. Transferrin endocytic trafficking followed in N2a cells transiently expressing MYC (control), BIN1-WT, BIN1-PL, and -KR, using a pulse-chase assay with fluorescently labeled transferrin (Alexa647-transferrin), analyzed by epifluorescence microscopy. **(A)** Endocytosed transferrin (2 min Alexa647-transferrin, fire LUT) detected in N2a cells. Insets show MYC signal corresponding to MYC (control), BIN1-WT, BIN1-PL, and -KR. Scale bar, 10 μm. **(B)** Quantification of endocytosed transferrin mean fluorescence in the percentage of control (n=3, N_control_=109 cells, N_Bin1-WT_=105 cells, N_Bin1-PL_=101 cells, N_Bin1-KR_=111 cells; \*\*\*\**P*<0.0001 Bin1-WT vs. control, Bin1-PL vs. control, Bin1-KR vs. control, Kruskal-Wallis test, mean ± SD). **(C)** Non-recycled transferrin (fire) detected upon 10 min pulse with Alexa647-transferrin and 20 min chase in N2a cells. Insets show MYC signal corresponding to control, BIN1-WT, BIN1-PL, and -KR. Scale bar, 10 μm. **(D)** Quantification of non-recycled transferrin mean fluorescence in percentage of control (n=3, N_control_=76, N_Bin1-WT_=97, N_Bin1-PL_=88, N_Bin1-KR_=97; ^ns^*P*=0.8713 Bin1-WT vs. control, ^ns^*P*>0.9999 Bin1-KR vs. control, \**P*=0.0265 Bin1-WT vs. Bin1-PL, ***P=0.0002 Bin1-PL vs. control, ****P<0.0001 Bin1-PL vs. Bin1-KR, Kruskal-Wallis test, mean ± SD).

**Figure 5.**
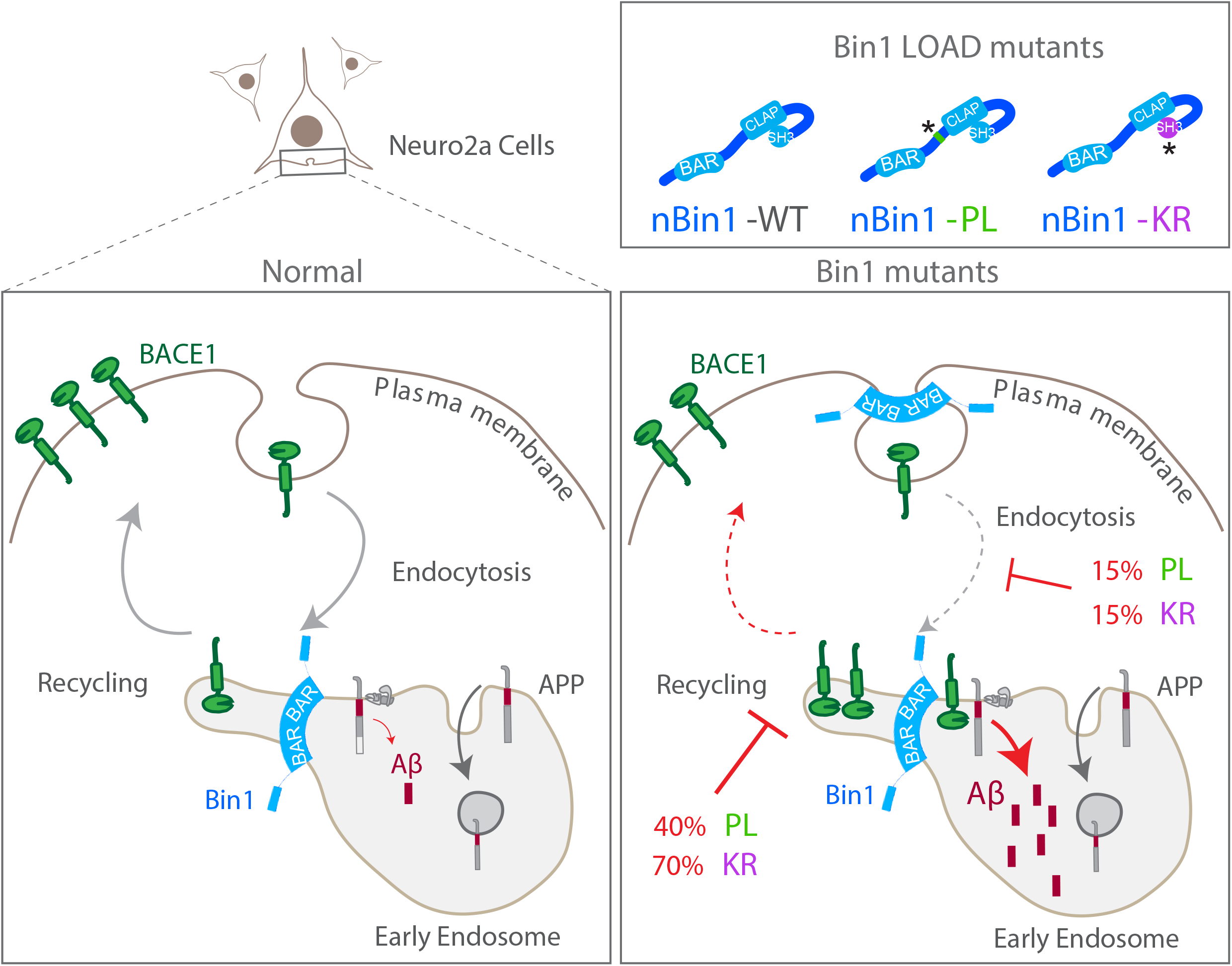
Schematic diagram illustrating the mechanisms used by Bin1 mutants to increase intracellular Aβ accumulation. Normally, Bin1 enables BACE1 recycling to the plasma membrane maintaining low Aβ production. The LOAD mutations PL (near CLAP domain) and KR (in the SH3 domain) in Bin1 interfere with its function in regulating BACE1 trafficking. Bin1 mutants interfere with BACE1 endocytosis but even more with its recycling—the accumulation of non-recycled BACE1 results in more Aβ production and accumulation. KR mutation leads to a more prominent defect in BACE1 recycling than the PL mutation. Thus, their pathogenicity may impact the development of late-onset AD early mechanisms differently.

Secondly, we measured non-recycled BACE1 (Fig. 3F and G). We found that non-recycled BACE1 was significantly increased in cells overexpressing Bin1-PL (43%) and Bin1-KR (70%) while Bin1-WT overexpression did not alter non-recycled BACE1 as compared to control (Fig. 3F).

The non-recycled BACE1 results from the net balance between endocytosis and recycling. Since BACE1 endocytosis decreased upon Bin1 mutant’s overexpression, the increase in non-recycled BACE1 is likely due to the reduction in BACE1 recycling.

We showed that Bin1 KD decreases BACE1 recycling resulting in BACE1 accumulation in early endosomes (15), and Miyagawa et al. showed BACE1 accumulation in late-endosomes/lysosomes overexpression. Overall non-recycled BACE1 colocalized more with EEA1-positive endosomes (40-(50). Thus, we investigated the localization of non-recycled BACE1 upon Bin1 mutants’ 50%) than with LAMP1-positive endosomes (10-15%). Notably, Bin1-PL and Bin1-KR overexpression mutants increased colocalization with LAMP1 (Fig. 3J and K). Bin1-WT overexpression did not affect increased non-recycled BACE1 colocalization with EEA1 by 12% (Fig. 3H and I). None of the Bin1 non-recycled BACE1 colocalization with EEA1 but increased colocalization with LAMP1 by 5% (Fig. 3H-K).

Together, our results indicate that Bin1-WT OE may not alter Aβ42 accumulation because it reduces BACE1 endocytosis required for APP cleavage by BACE1 or because it increases traffic to late-endosomes. In turn, both Bin1 mutants’ dominant-negative impact on BACE recycling was more prominent than on its endocytosis. Consequently, the decrease of BACE1 recycling to the plasma membrane leads to BACE1 accumulation in early endosomes, where likely increases APP processing. The mechanism may be due to the reduced interaction of Bin1 with BACE1 due to LOAD mutations.

### Bin1 mutants impair the canonical transferrin endocytic recycling

Since transferrin is the canonical cargo of endocytic recycling and Bin1 overexpression impairs transferrin endocytosis in non-neuronal cells (38), we next checked whether overexpression of Bin1-WT and mutants alter transferrin endocytosis and recycling, similarly to BACE1 (Fig 3).

To follow transferrin endocytosis, we pulsed N2a cells with fluorophore-conjugated transferrin for 2 min, upon Bin1-WT, Bin1-PL, and Bin1-KR OE. We found that Bin1-WT OE reduced transferrin internalization by 25% and that when cells overexpressed Bin1-PL and Bin1-KR, there was a similar reduction in transferrin endocytosis (20%) (Fig. 4A and B). These results indicate that Bin1-WT overexpression impairs transferrin endocytosis, which is not altered by the mutations.

To follow transferrin recycling, we chased endocytosed transferrin for 20 min after a 10 min pulse and quantified the intensity remaining intracellularly (Fig. 4C and D). Like BACE1, Bin1-WT OE did not significantly change while the Bin1-PL OE increased substantially by 50% non-recycled transferrin. Unexpectedly, the Bin1-KR OE did not alter the non-recycled transferrin.

These data indicate that transferrin recycling is reduced by Bin1 PL mutant but not by Bin1-WT or Bin1-KR. The KR mutation may have a dominant-negative effect more specific for the endocytic recycling of BACE1, while the PL mutation may have a more general dominant-negative effect on endocytic recycling.

## Discussion

BIN1 variants were associated with LOAD, but their translation into a disease mechanism is missing. Previously, we showed that Bin1 loss of function induced by Bin1 knockdown (KD) increased Aβ42 production due to the accumulation of BACE1 at enlarged early endosomes. Mechanistically, we found that Bin1 KD reduced BACE1 recycling to the plasma membrane. Here we investigated the impact of two LOAD coding variants in BIN1 function in controlling Aβ42 production and early endosome size. We introduced the two mutations, PL and KR, in wild-type neuronal BIN1 cDNA and determined the impact of their overexpression or re-expression in neuronal cells. Bin1 mutants increased Aβ42 intracellular accumulation, failed to control early endosome size, decreased BACE1 recycling, and reduced interaction with BACE1, indicating that the two LOAD mutations are pathogenic.

### Intracellular Aβ

The overexpression of Bin1 mutants increased intracellular Aβ42 while, in contrast, Bin1 wild-type did not. The Bin1 mutants’ phenotype was similar to that of Bin1 KD, although the mutants’ effect was more modest, increasing Aβ42 by 30%, while knocking down Bin1 increased Aβ42 by 65%. The loss of Bin1 function due to the LOAD mutations was confirmed when both mutants’ re-expression failed to rescue the Bin1 KD-dependent rise in Aβ42 compared to Bin1 wild-type. The impact of these two coding variants could be different from the regulatory variant (rs59335482) previously shown to increase BIN1 expression (42) since wild-type neuronal Bin1 overexpression did not impact intracellular Aβ42 accumulation. Since the increased expression of BIN1 also leads to increase levels of the ubiquitous Bin1 isoform, it remains undetermined if ubiquitous Bin1 overexpression affects intracellular Aβ42.

Interestingly, we also observed that the Bin1 mutants did not increase iAβ40, not recapitulating our previous observations that Bin1 KD increased iAβ40 although to a lesser extent than iAβ42 (15). Our results suggest an increase in the ratio Aβ42/40 in the presence of LOAD Bin1 mutations. The increase in the ratio Aβ42/40 in AD was established due to familial AD mutations in presenilins, the catalytic component of γ-secretase(61). More recent work indicates that presenilins’ activity depends on the trafficking of γ-secretase to endosomes and endosomal luminal pH (18, 62, 63). Thus the trafficking of γ-secretase may also be susceptible to the interference of LOAD mutations in the Bin1 control of endocytic recycling.

### Early endosome size

The Bin1 wild-type overexpression reduced Rab5 and EEA1-positive early endosome size as previously reported for Rab5-positive endosomes (51). The overexpression of Bin1 mutants did not recapitulate Bin1 wild-type reduction of early endosome size, suggestive of a loss of function. However, their loss of function was insufficient to induce early endosomes’ enlargement as observed when Bin1 is depleted. Interestingly, the rescue experiments revealed that the KR mutation in Bin1 could be more pathogenic than the PL mutation. Since Bin1-KR re-expression could not rescue Rab5-positive endosome size in cells depleted for Bin1, while the Bin1-PL re-expression partially rescued.

### BACE1 endocytic recycling

Given the previously reported impact of Bin1 overexpression in transferrin endocytosis (38, 64), we analyzed whether Bin1 wild-type and mutants’ overexpression impact transferrin and BACE1 endocytosis in neuronal cells. Our data confirmed that Bin1, when overexpressed, interferes with endocytosis, giving a similar inhibition for transferrin and BACE1 endocytosis. However, unexpectedly, the mutations in Bin1 did not alter the decrease in endocytosis induced by Bin1 overexpression. The mechanism of BACE1 endocytosis is somewhat controversial, with reports supporting that it is clathrin-mediated, while others indicate that it is ARF-6 dependent (7, 8). Our results support that BACE1 undergoes clathrin-mediated endocytosis like transferrin (65).

The overexpression of both Bin1 mutants reduced BACE1 recycling, recapitulating the Bin1 KD phenotype, indicating that the mutations have a dominant-negative effect inducing the loss of function of Bin1 in the control of BACE1 recycling. Accordingly, both mutations decreased Bin1 interaction with BACE1. The decrease in recycling leads to increased intracellular BACE1 in early endosomes, suggesting increased BACE1 access to APP in early endosomes, processing it more. We analyzed APP processing upon Bin1 mutants’ overexpression, but we found no significant change in the overall APP C-terminal fragments. The increase in beta-CTFs production is likely below the detection limit of the technique used. Unfortunately, more sensitive methods are not available for mouse APP-CTFs.

Nevertheless, we detected an increase in Aβ42 accumulation in early endosomes and late endosomes/lysosomes. Interestingly, the impact of the two mutations in Bin1 in BACE1 recycling has different magnitudes. The KR mutation has a more significant effect than the PL mutation in BACE1 recycling, which correlates with higher intracellular Aβ42.

Besides, we analyzed transferrin recycling since it decreases upon Bin1 depletion (40, 41) to determine if it was similarly affected by Bin1 mutants. Surprisingly we found that the PL mutation, but not KR, leads to decreased transferrin recycling. Thus, PL mutation likely interferes with Bin1 function in the control of endocytic recycling differently from KR mutation.

Overall, we find that the more prominent defect in BACE1 recycling plausibly overcomes the reduction in endocytosis to underlie the increase in Aβ42 production induced by mutant Bin1.

### Potential mechanisms

#### Bin1 WT overexpression impact in reducing endocytosis

Bin1 overexpression may inhibit endocytosis by sequestering its interacting partners’ clathrin, endophilin, and dynamin, indirectly compromising endocytic vesicle formation, membrane curvature, and scission, respectively (38, 64).

#### Bin1 mutants’ impact in reducing endocytic recycling

The difference in the PL and KR Bin1 mutants’ impact on endocytic recycling could be related to how the two mutations alter the Bin1 protein.

The PL mutation localizes in the proline-serine rich domain, where the lost proline could lead to an altered Bin1 secondary structure or conformation (66) or the observed reduced binding to BACE1 directly or via interactors (67), such as clathrin (68), which can participate in endocytic recycling (69).

The KR mutation occurs in the SH3 domain in the RT loop, one of three loops that characterize the structure of SH3 domains, composed of a patch of aromatic residues on the domain’s ligand-binding face that can modulate binding typically to proline-rich domains of proteins. Indeed, Bin1 interacts through its SH3 domain with the proline-rich domain in dynamin (70, 71). Further, the replacement of arginine to lysine was observed to reduce protein binding affinity due to conformation changes (67). Indeed, the KR mutation had a more dramatic impact in the interaction with BACE1, but the Bin1-KR interaction with dynamin, and others, maybe equally reduced.

Both mutations may also interfere with neuronal Bin1 self-inhibition through the intramolecular binding between the CLAP domain and the SH3 domain (47). This intramolecular binding may function as a curvature sensor triggering Bin1 to engage with the appropriately curved membrane (72). Thus, mutations in each domain could interfere with the Bin1 localization to the tubulated recycling endosomal carriers. Alternatively, it could interfere with the binding of Bin1 directly to BACE1 or indirectly through interaction with sorting nexin 4 (SNX4) (73), which also plays a role in BACE1 (74, 75) and transferrin recycling (76, 77).

Further studies of Bin1 mutations are underway, namely identifying Bin1 mutants interactome, which should illuminate the mechanism of interfering with BACE1 recycling.

### Bin1 as an AD risk factor

These results support the known function of Bin1 in BACE1 recycling at early endosomes and show how Bin1 variants associated with LOAD interfere with this function, potentiating Aβ42 production and intracellular accumulation as well as endosome enlargement. The mild loss of Bin1 PL and KR mutants’ function is consistent with developing a late-onset form of AD. Future studies are underway to confirm if this intracellular accumulation of Aβ42 is sufficient to cause synaptic dysfunction, the most critical effector of AD cognitive decline.

## EXPERIMENTAL PROCEDURES

### cDNA and siRNA

We used the following DNA plasmids encoding: BACE1-GFP (15); FLAG-BACE1-GFP (15); Rab5-GFP plasmid was a gift from M. Arpin (Institut Curie); siRNA-resistant neuronal Bin1-MYC construct (brain amphiphysin II (BRAMP2); isoform 1; NP_033798.1; (15)); Bin1-PL and Bin1-KR were generated by site-directed mutagenesis (NZYtech) of siRNA-resistant neuronal Bin1-MYC (for Bin1-PL, primers 5’GAACCATGAGCCAGAGCTGGCCAGTGGGGCCTC’ and 5’GAGGCCCCACTGGCCAGCTCTGGCTCAT GGTTC’; for Bin1-KR primers 5’GATGAGCTGCAACTCAGAGCTGGCGATGTGGTG’ and 5’CACCACATCGCCAGCTCTGAGTTGCAGCTCATC3). All plasmids were sequenced. We used the following siRNA oligonucleotides: as siControl a non-targeting control siRNA (GeneCust) and for Bin1 knockdown, siBin1 (65598; ThermoFisher Scientific) (15).

### Cell culture, transfections and treatments

Neuroblastoma Neuro2a (N2a) cells (ATCC®CCL-131™) were a gift from Zsolt Lenkei (ESPCI-ParisTech). Cells were cultured in DMEM-GlutaMAX (ThermoFisher Scientific) with 10% FBS (Sigma-Aldrich) at 37°C in 5% CO_2_. For cDNA expression, N2a cells were transiently transfected with 0.5 ug of cDNA with Lipofectamine 2000 (ThermoFisher Scientific) and analyzed after 24h. For small interfering RNA (siRNA) treatment, N2a cells were transiently transfected with 10 nM specific siRNA with Lipofectamine RNAiMax (ThermoFisher Scientific) and analyzed after 72 h. When indicated, cDNA was transfected after 48 h of siRNA treatment, and cells were analyzed after 24 h. When indicated, BACE1 was inhibited by 9 to 16 h treatment with 30 µM compound IV (Calbiochem), gamma-secretase was inhibited by 9 to 16 h treatment with 250 nM DAPT (Calbiochem), or DMSO (solvent) was used as control. All experiments were carried out in at least three independent sets of culture, except when indicated.

### Antibodies and probes

The following antibodies were used: anti-APP (Y188, GeneTex, cat GTX61201, 1:1,000); anti-Aβ42 mAb (H31L21, Invitrogen, cat 700254, 1:200); anti-Aβ40 pAb (Sigma, cat Ab5074P, 1:100); anti-FLAG (M1) mAb (Sigma, cat F3040, 1:100); anti-MYC pAb (1:500); anti-tubulin mAb (Tu-20, Millipore, cat MAB1637, 1:10,000); anti-EEA1 pAb (Sigma, cat E3906; 1:250); anti-EEA1 pAb (N-19, Abcam, cat sc-6415, 1:50); anti-LAMP1 mAb (BD, cat 553792, 1:200). The probe Alexa Fluor™ 647 Conjugate-transferrin (ThermoFisher Scientific) was used in pulse-chase assays. Immunofluorescence labeling N2a cells were fixed with 4% paraformaldehyde for 10 to 15 min, permeabilized and blocked with 0.1% saponin, 2% FBS, 1% BSA for 1 h before antibody incubation using standard procedure. For Aβ42 labelling cells were permeabilized with 0,1% saponin for 1h before blocking, and primary antibody incubated for 16h. Coverslips were then mounted using Fluoromount-G (Southern Biotech).

### Trafficking assays

Trafficking assays were performed as previously (15). Briefly, before endocytosis experiments, N2a cells, expressing FLAG-BACE1-GFP or not, were starved in a serum-free medium (30 min). For BACE1 endocytosis, N2a cells were incubated with anti-FLAG (M1) for 10 min. For BACE1 recycling, N2a cells pulsed with M1 were acid stripped (0.5 M NaCl, 0.2 M acetic acid; 4 s) and quickly rinsed in PBS before chasing for 20 min at 37°C. A second acid stripping was performed before fixation, and non-recycled proteins were immunolabelled upon permeabilization. For transferrin endocytosis, N2a cells were incubated with fluorophore-conjugated transferrin for 2 min. For transferrin recycling, N2a cells pulsed with fluorophore-conjugated transferrin were washed in PBS (4 s) before recycling for 20 min at 37°C. A second washing was performed before fixation.

### Image acquisition

Epifluorescence microscopy was carried out on a widefield upright microscope Axio Imager.Z2 (Zeiss, Oberkochen, Germany) equipped with a 60×NA-1.4 oil immersion objective and an AxioCam MRm CCD camera (Zeiss). Confocal microscopy was performed with LSM980 equipped with AiryScan 2.0 (Zeiss). For direct comparison, samples were imaged in parallel and using identical acquisition parameters.

### Image processing and analysis

Image processing analyses were carried out using the free image analysis software: ICY (78) and FIJI (ImageJ) (79).

For the quantification of intracellular Aβ42, BACE1, and transferrin levels, regions of interest corresponding to randomly chosen single cells were outlined based on MYC labeling using the ICY “polygon type ROI (2D” tool. An ROI was also randomly chosen in an area without cells corresponding to the background. The mean fluorescence in each ROI was obtained using ICY ROI export and was presented as a percentage of the indicated control upon background fluorescence subtraction.

For the quantification of Rab5- and EEA1-positive endosomes puncta size and density per area, images were processed using “subtract background” in FIJI. Then the endosomes were segmented using the ICY “Spot detector” plugin in each single cell ROI. The number of spots per cell ROI area was used to obtain the endosome density per area.

For the line profiles of Aβ, Bin1 and MYC we used the “plot profile” tool in FIJI.

For the quantification of colocalizations we used the ComDet v.0.5.5 plugin for Image J *(https://github.com/ekatrukha/ComDet)*.”

### Immunoblotting

N2a cell lysates were prepared using modified RIPA buffer (50 mM Tris–HCl pH 7.4, 1% NP-40, 0.25% sodium deoxycholate, 150 mM NaCl, 1 mM EGTA, 1% SDS, with 1X PIC (Sigma-Aldrich). Sonication was performed with the settings: 3 cycles of 1s on and 45ms off (pulse; total time of 30 s) at 10% amplitude. Proteins separated by 7.5, 10, or 4-12% Tris-glycine SDS–PAGE was transferred to 0.45 μm nitrocellulose membranes and processed for standard immunoblotting. HRP-conjugated secondary antibodies signal was revealed using ECL Prime kit (GE Healthcare) and captured using the ChemiDoc imager (BioRad) within the linear range and quantified by densitometry using the ImageJ software protocol (https://imagej.nih.gov/ij/docs/menus/analyze.html#gels).

### Co-immunoprecipitation

Adult wild-type mice (BALB/c) cortices lysates were immunoprecipitated with anti-Bin1 or mouse IgG 16 h at 4 °C and then with 30 μl of protein G–Sepharose beads (GE Healthcare) for 3 h at 4°C. Beads were washed 3x with lysis buffer. The sample was eluted with 2x SDS loading buffer, resolved by 7.5% Tris-Glycine SDS-PAGE and detected by immunoblotting. Animal procedures were performed according to EU recommendations and approved by the NMS-UNL ethical committee (07/2013/CEFCM), and the national DGAV (0421/000/000/2013).

N2a cell lysates were prepared using modified RIPA buffer (50 mM Tris–HCl pH 7.4, 1% NP-40, 0.25% sodium deoxycholate, 150 mM NaCl, 1 mM EGTA, 1% SDS, with 1X PIC (Sigma-Aldrich). Lysates were immunoprecipitated with GFP-Trap® Agarose beads (Chromotek) following the manufacturer protocol. Briefly, lysates were rotated with GFP-Trap® Agarose beads for 1 h at 4°C. Beads were washed three times with washing buffer (10 mM Tris-Cl pH 7.5; 100 mM NaCl; 0.5 mM EDTA; 0.05% NP-40; 1% glycerol). The IP proteins were eluted with 2x SDS loading buffer, resolved by 7.5% Tris-Glycine SDS-PAGE together with lysates (input) and detected by immunoblotting.

### Statistics

GraphPad Prism 8 software was used for graph generation with mean ± SD. The sample size was determined based on pilot studies. Statistical significance for at least three independent experiments was determined on normal data D’Agostino-Pearson omnibus normality test) by two-tailed Student’s t-test and multiple comparisons one-way ANOVA with Tukey’s test using GraphPad Prism 6. Statistical significance for nonparametric data was tested by the Mann–Whitney test or, for multiple comparisons, the Kruskal-Wallis test, followed by Dunn’s multiple comparison test. Data were expressed as mean ±SEM.

## Data availability

Data are to be shared upon request.

## Acknowledgments

We thank M. Arpin (I Curie) and Z. Lenkei (ESPCI-ParisTech) for the gift of plasmids and cells, respectively.

We thank Ana Cláudia Marques for her preliminary observations and lab members for helpful discussions and critical reading of the manuscript.

We thank S. Marques (CEDOC Animal Facility), T. Pereira (CEDOC Microscopy Facility), and Ana Oliveira (CEDOC Cell Culture Facility) for their technical assistance.

## Author contributions

C.G.A, C.B.P., M.B., and T.B. conceptualization; C.G.A data curation; C.G.A., C.B.P., and M.A.B. formal analysis; C.G.A validation; C.B.P, M.A.B, and T.B. investigation; M.A.B, C.B.P, and C.G.A visualization; C.G.A., C.B.P. M.A.B., and T.B. methodology; C.B.P writing-original draft; C.G.A., M.A.B. and C.B.P. writing-review and editing; C.G.A. supervision; C.G.A. funding acquisition; C.G.A project administration.

## Funding and additional information

This project has received institutional funding from iNOVA4Health— UID/Multi/04462/2019; H2020-WIDESPREAD-01-2016-2017-TeamingPhase2 - GA739572; the research infrastructure PPBI-POCI-01-0145-FEDER-022122, co-financed by FCT (Portugal) and Lisboa2020, under the PORTUGAL2020 agreement (European Regional Development Fund).

CGA has obtained funding from Maratona da Saúde 2016; CEECIND/00410/2017 financed by FCT (Portugal); ALZ AARG-19-618007 Alzheimer’s Association); the European Union’s Horizon 2020 research and innovation program under grant agreement No 811087 (Lysocil). CBP was the recipient of an FCT doctoral fellowship (PD/BD/128374/2017). TB has been the recipient of an FCT doctoral fellowship (SFRH/BD/131513/2017). MB is the recipient of an FCT doctoral fellowship (2020.06758.BD).

## Abbreviations

AD: Alzheimer’s disease
AP-2: adaptor protein complex 2
APOE4: Apolipoprotein E ε4
APP: amyloid precursor protein
ARF6: ADP-ribosylation factor 6
Aβ: amyloid-beta
BACE1 β-site: APP-cleaving enzyme 1 or β-secretase 1
BAR: BIN1/amphiphysin/RVS167 domain
BIN1: Bridging integrator 1 / MYC box-dependent-interacting protein 1
BSA: bovine serum albumin
CTF: carboxyl-terminal fragment
cDNA: complementary DNA
CLAP: clathrin and AP2 binding domain
DMEM: Dulbecco’s Modified Eagle Medium
EEA1: Early endosome antigen 1
FAD: familial Alzheimer’s disease
EGTA: Ethylene Glycol Tetra-acetic Acid
FBS: fetal bovine serum
GFP: Green Fluorescent Protein
GWAS: genome-wide association studies
HRP: Horseradish peroxidase
IP: Immunoprecipitation
IgG: Immunoglobulin G
KD: Knockdown
LOAD: late-onset Alzheimer’s disease
N2a: Neuro2a
mAb: monoclonal antibody
OE: overexpression
pAb: polyclonal antibody
PBS: phosphate-buffered saline
PIC: protease Inhibitor Cocktail
RIPA: Radio-Immunoprecipitation Assay
RVS167: Reduced viability upon starvation protein 167
SDS-PAGE: sodium dodecyl sulfate-polyacrylamide gel electrophoresis
SH3: src homology 3 domain
siRNA: small interfering RNA
SNX4: sorting nexin-4

## Figure legends

**Figure S1.**
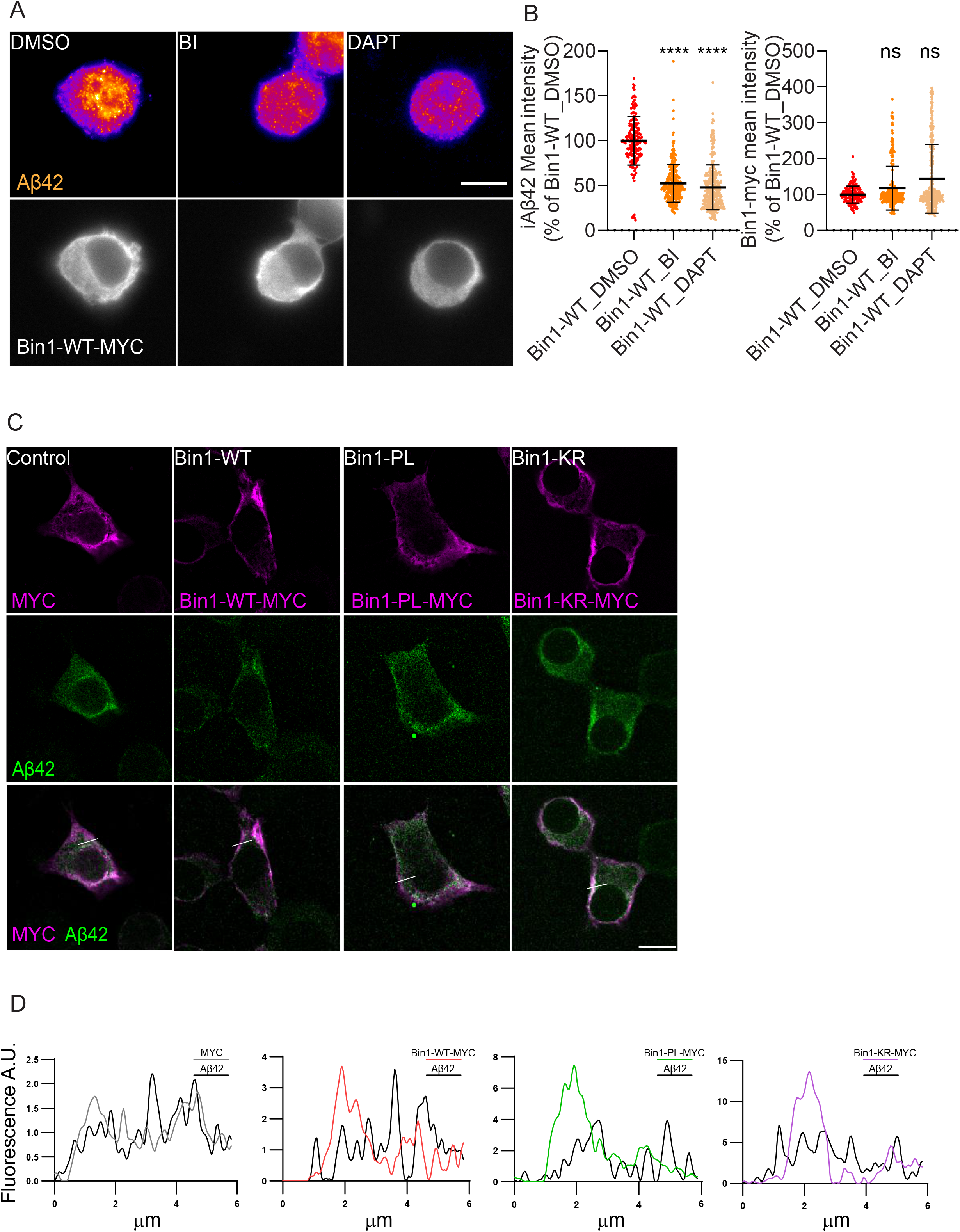
Aβ42 immunofluorescence is specific and independent of MYC signal. **(A)** Intracellular endogenous Aβ42 (fire LUT) in N2a cells transiently expressing Bin1 wild-type (WT) treated with β-secretase inhibitor IV (BI), DAPT or DMSO (control), immunolabelled with anti-Aβ42 (upper panels) and anti-MYC (lower panels), analyzed by epifluorescence microscopy. Scale bar, 10 μm. **(B)** Quantification of Aβ42 and MYC fluorescence mean intensity (n=3, N_BIN1-WT_DMSO_=194 cells, N_Bin1-WT_BI_=289 cells, N_Bin1-WT_DAPT_=344 cells; ^ns^*p*>0.9999 Bin1-WT_BI vs. Bin1-WT_DMSO, ^ns^*p*=0.2474 Bin1-WT_DAPT vs. Bin1-WT_DMSO, \*\*\*\**P*<0.0001 Bin1-WT_BI vs. Bin1-WT_DMSO, Bin1-WT_DAPT vs. Bin1-WT_DMSO, Kruskal-Wallis test, mean ± SD). **(C)** High resolution imaging (airyscan) of intracellular endogenous Aβ42 in N2a cells transiently expressing Bin1-WT, Bin1-PL and Bin1-KR tagged with MYC, or MYC (control), immunolabelled with anti-MYC (magenta, upper panels) and anti-Aβ42 (green, middle panels). Merged imaged (lower panels) analyzed by confocal microscopy. Scale bar, 10 µm **(D)** Line intensity profiles of Aβ42 (black line) and MYC indicated in (C), in N2a cells transiently expressing MYC (grey line), Bin1-WT-MYC (red line), Bin1-PL-MYC (green line) and Bin1-KR-MYC (purple line).

**Figure S2.**
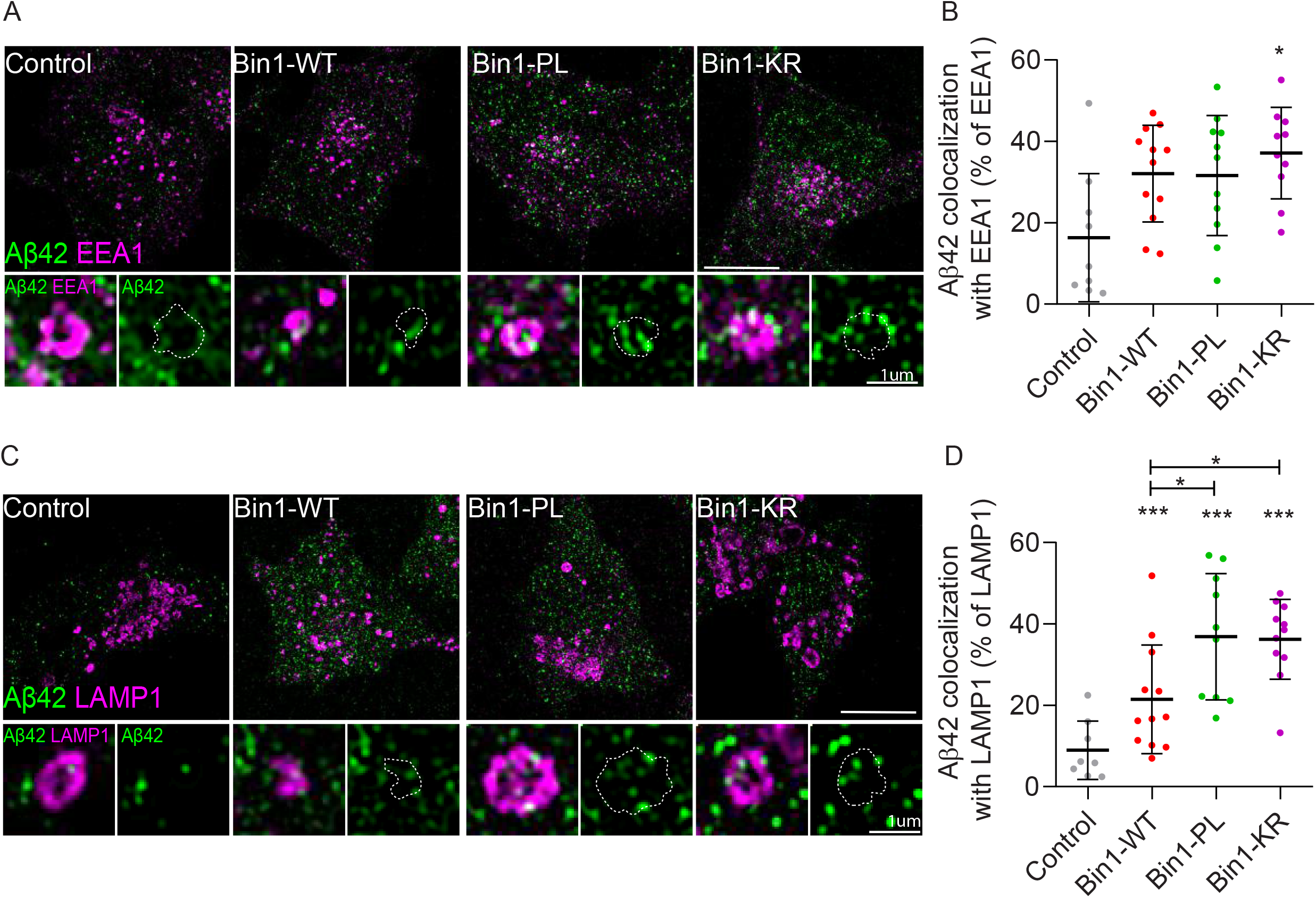
Aβ42 accumulation in early and late-endosomes. **(A)** Intracellular endogenous Aβ42 (green) localization in EEA1-positive endosomes (magenta) in N2a cells transiently expressing Bin1-WT, Bin1-PL, Bin1-KR tagged with MYC, or MYC (control), immunolabelled with anti-Aβ42 and anti-EEA1, and analyzed by Airyscan confocal microscopy. Images are displayed merged. Scale bar, 10 µm. Insets show magnified endosomes, Aβ42 is shown merged with EEA1 and individually. Scale bar, 1 µm. **(B)** Quantification of EEA1 colocalization with Aβ42 (n=2, Ncontrol=9 cells, NBin1-WT=12 cells, NBin1-PL=11 cells, NBin1-KR=10 cells; nsP=0.1825 Bin1-WT vs. control, Bin1-PL vs. control, *P=0.0346 Bin1-KR vs. control, Kruskal-Wallis test, mean ± SD). **(C)** Intracellular endogenous Aβ42 (green) localization in LAMP1-positive endosomes (magenta) in N2a cells transiently expressing Bin1-WT, Bin1-PL, Bin1-KR tagged with MYC, or MYC (control), immunolabelled with anti-Aβ42 and anti-LAMP1 and analyzed by Airyscan confocal microscopy. Images are displayed merged. Scale bar, 10 µm. Insets show magnified endosomes, Aβ42 is shown merged with LAMP1 and individually. Scale bar, 1 µm. **(D)** Quantification of LAMP1 colocalization with Aβ42 (n=2, N_control_=8 cells, N_Bin1-WT_=12 cells, N_Bin1-PL_=10 cells, N_Bin1-KR_=11 cells; ^ns^*P=*0.1267 Bin1-WT vs. control, *P=0.0262 Bin1-WT vs. Bin1-PL; \**P*=0.0293 Bin1-WT vs. Bin1-KR, \*\*\**p*=0.0001 Bin1-KR vs. control, Bin1-PL vs. control, Kruskal-Wallis test, mean ± SD).

**Figure S3.**
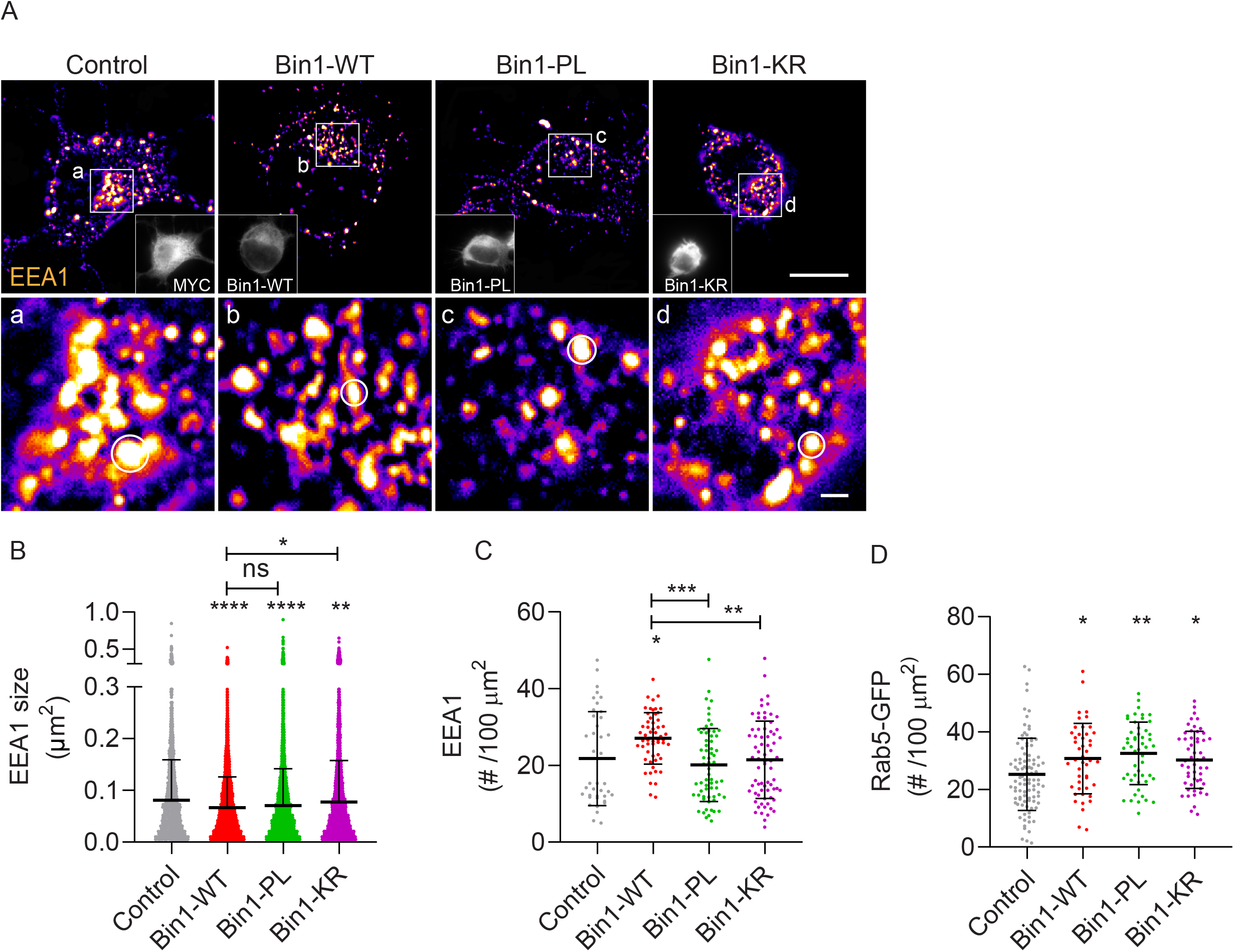
Bin1mutants’ impact EEA1 endosome size. **(A)** EEA1-positive early endosomes (fire LUT) detected by immunofluorescence with an antibody against EEA1 in N2a cells transiently expressing MYC (control), BIN1-WT, BIN1-PL, and -KR tagged with MYC (insets), analyzed by epifluorescence microscopy. Scale bar, 10 μm. The white squares indicate the perinuclear region magnified in (a-d). Circles highlight the mean size of EEA1-positive early endosomes. Scale bar, 1 μm. **(B)** Quantification of EEA1-positive endosomes size (µm^2^) (n=2, N_control_=39 cells, N_Bin1-WT_=55 cells, N_Bin1-PL_=65 cells, N_Bin1-KR_=72 cells; ^ns^*P*>0.9999 Bin1-WT vs. Bin1-PL, \**P*=0.0415 Bin1-WT vs. Bin1-KR, \*\**P*=0.0070 Bin1-KR vs. control, \*\*\*\**P*<0.0001 Bin1-WT vs. control, Bin1-PL vs. Control, Kruskal-Wallis test, mean ± SD). **(C)** Quantification of EEA1-positive endosomes number per 100 µm^2^ (n=2, N_control_=39 cells, N_Bin1-WT_=55 cells, N_Bin1-PL_=65 cells, N_Bin1-KR_=72 cells; \**P*=0.0288 Bin1-WT vs. control, \*\**P*=0.0052 Bin1-WT vs. Bin1-KR, \*\*\**P*=0.0004 Bin1-WT vs. Bin1-PL, Kruskal-Wallis test, mean ± SD). **(D)** Quantification of Rab5-positive endosomes number per 100 µm^2^ (n=3, N_control_=96 cells, N_Bin1-WT_=45 cells, N_Bin1-PL_=51 cells, N_Bin1-KR_=53 cells; \**P*=0.0451 Bin1-WT vs. control, \**P*=0.0393 Bin1-KR vs. control, \*\**P*=0.0013 Bin1-PL vs. control, Kruskal-Wallis test, mean ± SD).

**Figure S4.**
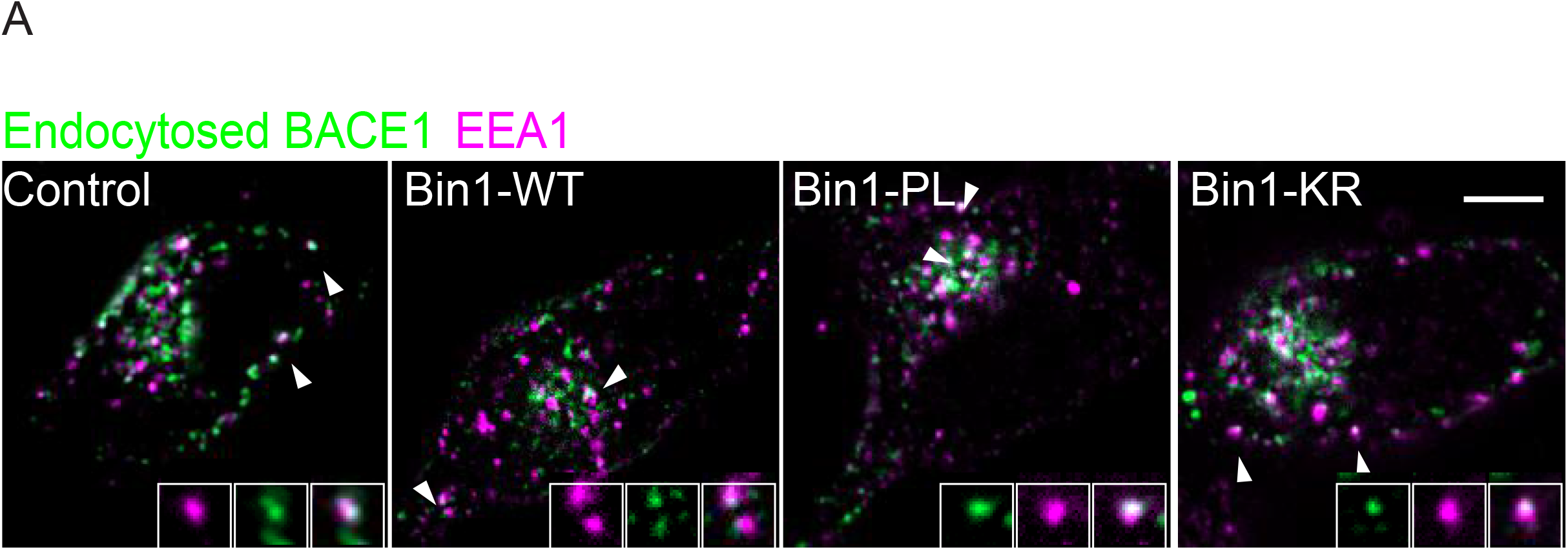
Endocytosed BACE1 colocalizes with EEA1. (**A**) Endocytosed BACE1 (green, detected upon 10 min pulse with M1) localization in EEA1-positive endosomes (magenta), assessed by immunofluorescence in N2a cells with an antibody against endogenous EEA1. Images are displayed merged after background subtraction with Fiji. Scale bar, 10 μm. The white squares indicate magnified endosomes.

